# Acute phase protein, α – 1- acid glycoprotein (AGP-1), has differential effects on TLR-2 and TLR-4 mediated responses

**DOI:** 10.1101/634295

**Authors:** Mosale Seetharam Sumanth, Kandahalli Venkataranganayaka Abhilasha, Shancy Petsel Jacob, Vyala Hanumanthareddy Chaithra, Venkatesha Basrur, Belinda Willard, Thomas M. McIntyre, K. Sandeep Prabhu, Gopal K. Marathe

## Abstract

Alpha-1-acid glycoprotein (AGP-1) is a major positive acute phase glycoprotein with unknown functions that likely plays a role in inflammation. We tested its involvement in a variety of inflammatory responses using human AGP-1 purified to apparent homogeneity and confirmed its identity by immunoblotting and mass spectrometry. AGP-1 alone upregulated MAPK signaling in murine peritoneal macrophages. However, when given in combination with TLR ligands, AGP-1 selectively augmented MAPK activation induced by ligands of TLR-2 (Braun lipoprotein) but not TLR-4 (lipopolysaccharide). *In vivo* treatment of AGP-1 in a murine model of sepsis with or without TLR-2 or TLR-4 ligands, selectively potentiated TLR-2-mediated mortality, but was without significant effect on TLR-4-mediated mortality. Furthermore, *in vitro*, AGP-1 selectively potentiated TLR-2 mediated adhesion of human primary immune cell, neutrophils. Hence, our studies highlight a new role for the acute phase protein AGP-1 in sepsis via its interaction with TLR-2 signaling mechanisms to selectively promote responsiveness to one of the two major gram-negative endotoxins, contributing to the complicated pathobiology of sepsis.

## Introduction

Sepsis is a dysregulated inflammatory disorder that follows infection and is a major leading cause of mortality globally in the intensive care units (ICU) (Singer, et al., 2016, Chaudhry and Duggal, 2014, Phua, et al., 2013). Bacterial endotoxins such as the lipopolysaccharide (LPS) or membrane lipoproteins such as Braun lipoprotein (BLP) have often been used to mimic sepsis in *in vivo* models (Lakshmikanth, et al., 2016, Remick, et al., 2000, Jacob, et al., 2016, Lakshmikanth, et al., 2016). Sepsis involves a complex interplay of proinflammatory and anti-inflammatory actions in response to multiple mediators (Frost, et al., 2002, Lu, et al., 2008). LPS and BLP are detected by the pattern recognition receptors (PRRs), TLR-4 and TLR-2, respectively (Lakshmikanth, et al., 2016, Maeshima and Fernandez, 2013), to upregulate myriad of genes that amplify the inflammatory response (Lu, et al., 2008). The initial hyper-inflammatory response during sepsis is counterregulated by the compensatory anti-inflammatory response (CAR) (van der Poll and van Deventer, 1999). Endogenous serum factors including acute phase proteins are part of CAR and help in maintaining homeostasis (Johnson, et al., 1999). Alpha–1–acid glycoprotein (AGP-1) is a major acute phase glycoprotein, whose levels increase several folds during sepsis (Fournier, et al., 2000). AGP-1, a positive acute phase protein, has an unusually low pI of 2.8 – 3.8 and contains glycosyl moieties accounting for up to 45% of its mass (Fournier, et al., 2000, Schmid, et al., 1980). Numerous biological activities have been attributed to AGP-1 (Hochepied, et al., 2003), yet its exact function(s) remain to be delineated. Both elevated (Frazier, 2011) and decreased (Barroso-Sousa, et al., 2013) levels of AGP-1 have been reported in sepsis, with correlation to mortality. In isolation, AGP-1 inhibits neutrophil migration to the infection site via nitric oxide, promoting inadequate pathogen clearance (Mestriner, et al., 2007). On the contrary, increased AGP-1 concentration is reported to limit adverse inflammatory reactions through a negative feedback loop (Libert, et al., 1994, Roitt and Delves, 1998).

This study determined whether AGP-1 modulates endotoxin-mediated sepsis using isolated cells and an *in vivo* murine model of sepsis (Lakshmikanth, et al., 2016). We demonstrate that AGP-1 selectively augments lipoprotein-mediated inflammatory responses suggesting a novel mechanism of action.

## Materials and Methods

### Reagents

Cibacron F3GA agarose, DEAE-cellulose gel beads, anti-human AGP-1 antibody, anti-mouse-IgG-HRP antibody, commercial AGP-1, Foetal Bovine Serum (FBS), LPS and RPMI media were procured from Sigma Chemicals Co., St. Louis, MO, USA. Phospho-specific antibodies to p-38, JNK and ERK, β-actin and anti-rabbit IgG-HRP were obtained from Cell Signaling Technology, Danvers, MA, USA. Complete Mini EDTA-free protease inhibitor cocktail tablets were from Roche Diagnostics, Mannheim, Germany. PVDF membrane was from BioRad Laboratories, Hercules, California, USA and Thioglycollate media was obtained from Sisco Research Laboratories, Mumbai, India. Trypsin was bought from Promega, Madison, WI, USA. Calcein 2 AM and molecular weight markers were procured from Invitrogen, Carlsbad, CA, USA.

### Animals

Male and female Swiss albino mice, 8-10 weeks old, weighing 20-25g were obtained from Central Animal Facility, University of Mysore, and all the experiments were approved by Institutional Animal Ethics Committee, University of Mysore, Mysore, India (Approval No: UOM/IAEC/12/2016). The animals were housed with adequate ventilation, food, and water (available ad libitum) and were monitored daily over the experimental period.

### Serum collection

Blood was drawn from healthy human volunteers after obtaining informed consent. Permission to draw blood from human volunteers was also obtained from the Institutional Human Ethics Committee (UOM No. 104 Ph.D/2015-16). Briefly, the blood was collected and allowed to coagulate. The coagulated blood was then centrifuged at 600 x g for 20 min at 25□C to collect serum and stored at −20□C until further use.

### Purification of AGP-1 from human serum

Cross-linked agarose covalently attached to Cibacron blue F3GA dye was used as a simple first step in the purification of AGP-1 where serum proteins, including AGP-1 that do not bind to the gel, are eluted in the void volume (Caramelo-Nunes, et al., 2011). Briefly, human serum was loaded on to a Cibacron F3GA agarose column (30 x 1.5 cm) equilibrated with 10 mM phosphate buffer (pH 7.8). Unbound proteins containing AGP-1 were eluted in equilibration buffer at a flow rate of 1 ml/min as followed by UV absorbance at 280 nm. AGP-1 containing fractions were pooled and concentrated by spin concentrators (10 kDa cut off) (Agilent Technologies, USA). The concentrate was then applied to a DEAE-cellulose column (25 x 0.5 cm) equilibrated with 30 mM acetate buffer (pH 5.0). Fractions were eluted with buffer containing sodium chloride (0 – 2 M) at a flow rate of 24 ml/ hour. AGP-1 containing fractions were pooled, concentrated, and quantified using Lowry’s method (Lowry, et al., 1951). The endotoxin content of purified AGP-1 was assessed by Limulus amebocyte lysate (LAL) assay (Endochrome – K^TM^, Charles River, SC, USA) as per manufacturer’s instructions.

### SDS-PAGE and Western blotting for AGP-1

AGP-1 samples were resolved on a reducing SDS-PAGE (7.5 % acrylamide), visualized either by Coomassie brilliant blue or Periodic acid-Schiff (PAS) staining (Zacharius, 1969) and subjected to Western blotting. Blots were stained using appropriate primary antibody (anti-human AGP-1 monoclonal antibody; 1:10000 v/v) and secondary antibody (anti-mouse IgG HRP conjugate; 1:5000 v/v). The blots were visualized using freshly prepared ECL reagent by UV-transillumination (Uvi-Tech, Cambridge, UK).

### Mass spectral analysis of AGP-1

Purified samples were resolved on SDS-PAGE, visualized using Coomassie stain, AGP-1 excised, and the band destained with 30% methanol for 4 hours. Upon reduction (10 mM dithiothreitol) and alkylation (65 mM 2-chloroacetamide) of the cysteines, proteins were digested overnight with sequencing grade modified trypsin. The resulting peptides were resolved on a nano-capillary reverse phase column (Acclaim PepMap C18, 2 micron, 50 cm, Thermo Scientific) using a 1% acetic acid/acetonitrile gradient at 300 ml/min and directly introduced into Q Exactive HF mass spectrometer (Thermo Scientific, San Jose CA). MS1 scans were acquired at 60 K resolution (AGC target=3e6, max IT=50ms). Data-dependent high-energy C-trap dissociation MS/MS spectra were acquired for the 20 most abundant ions (Top20) following each MS1 scan (15 K resolution; AGC target=1e5; relative CE ~28%). Proteome Discoverer (V 2.1; Thermo Scientific) software suite was used to identify the peptides by searching the HCD data against an appropriate database. Search parameters included MS1 mass tolerance of 10 ppm and fragment tolerance of 0.1 Da. False discovery rate (FDR) was determined using Fixed PSM validator and proteins/peptides with a FDR of ≤1 % were retained for further analysis.

### Purification of Braun lipoprotein (BLP) from *E. coli* DH5α

Purification of BLP was carried out as before (Lakshmikanth, et al., 2015). The final precipitate was dissolved in 10 mM sodium phosphate buffer (pH 7.4) containing 1% SDS (S-buffer). The endotoxin content in the purified BLP was assessed by Limulus amebocyte lysate (LAL) assay using Endochrome – KTM kit (Charles River, SC, USA) as per manufacturer’s instructions.

### Isolation of murine peritoneal macrophages

Mice received an intraperitoneal injection of 3 ml thioglycollate broth (3% w/v). Macrophages were harvested 3 days later by washing peritoneal cavity with ice-cold PBS, and the resulting cells were resuspended in RPMI medium containing 2% FBS. These cells were incubated for 2 hours at 37°C and 5% CO_2_. Adherent cells were used as peritoneal macrophages.

### MAPK signaling

Adherent mouse peritoneal macrophages (2 x 10^6^ cells/ml) were stimulated for 30 min with AGP-1 (25, 50, and 100 μg/ml) with or without BLP (100 ng/ml) or LPS (100 ng/ml). Unstimulated macrophages were the negative control. Lysates were prepared using Radio-Immunoprecipitation Assay (RIPA) buffer and immunoblots were developed using specific primary and appropriate secondary antibodies for phospho-p38, phospho-JNK, phospho-ERK, and β-actin (1:1000 v/v).

### Murine endotoxemia

Murine endotoxemia was induced by administering the LPS *(E. coli* O111: B4) or BLP in the presence of the hepatotoxin D-galactosamine (Lakshmikanth, et al., 2016, Jacob, et al., 2016). BLP (5, 10, and 15 mg/kg) was administered intraperitoneally in the presence or absence of AGP-1 (25 mg/kg) to Swiss albino mice. LPS was injected intraperitoneally at 15 mg/kg in the presence or absence of AGP-1. Survival and clinical signs were monitored for 24 hours. All animals were euthanized 24 hours post-treatment with LPS or BLP.

### Neutrophil Adhesion Assay

Neutrophils were isolated freshly from a healthy human volunteer with informed consent. The neutrophils were isolated by dextran sedimentation and separation over ficoll density gradient centrifugation (Watanabe, et al., 2003). Neutrophil-rich pellet obtained was suspended in 1ml of Hank’s-Balanced salt solution (HBSS) containing 0.2% human serum albumin (HBSS/A). For assessment of adhesion, the neutrophil suspension was loaded with calcein-AM to a final concentration of 1 μ M prior to incubation for a period of 45 min at 37 ^0^C. The labeled neutrophils were incubated with LPS and BLP (100 ng/ml) with or without AGP-1 (25, 50, and 100 μg/ml) in triplicate wells in twelve well cell culture plates (Nest Biotechnology Co. Ltd., China) pre-coated with 0.2% gelatin. The adherent neutrophils were visualized and photographed at a magnification of 10x under a fluorescent microscope (Lakshmikanth, et al., 2016). The number of cells adhered in each well was determined by counting the cells adhered in 10 randomly chosen fields using ImageJ software and then calculating the average number of cells adhered per field.

### Statistical analysis

The animal experiments are representative of more than 2 independent experiments and the statistical significance among groups was determined by the log-rank test. Un-paired two-tailed t-test was used to compare the mean for each treatment group with the mean of the control group. All other results were analyzed using one-way analysis of variance (ANOVA) where applicable.

## Results

### Purification of AGP-1 from human serum and its identity with commercial AGP-1

Commercial purified human plasma AGP-1 showed batch-to-batch variation in the electrophoretic mobility of its components (data not shown), prompting us to purify AGP-1 in large quantities from healthy human volunteers. AGP-1 purified by conventional chromatographic techniques (Fig 1 A and B) showed apparent homogeneity (Figs 1 C, D, and E) as a 43 kDa glycoprotein. Further, LC/MS analyses of the purified tryptic peptides were compared with the commercially available AGP-1 and were found to be identical and mapped to the predicted human sequence. Isolated AGP-1 showed a protein mass of 23.5 kDa with a pI of 5.02 (Fig 1F). The endotoxin content of the purified AGP-1 was < 7 EU/mg protein.

**Figure 1:**
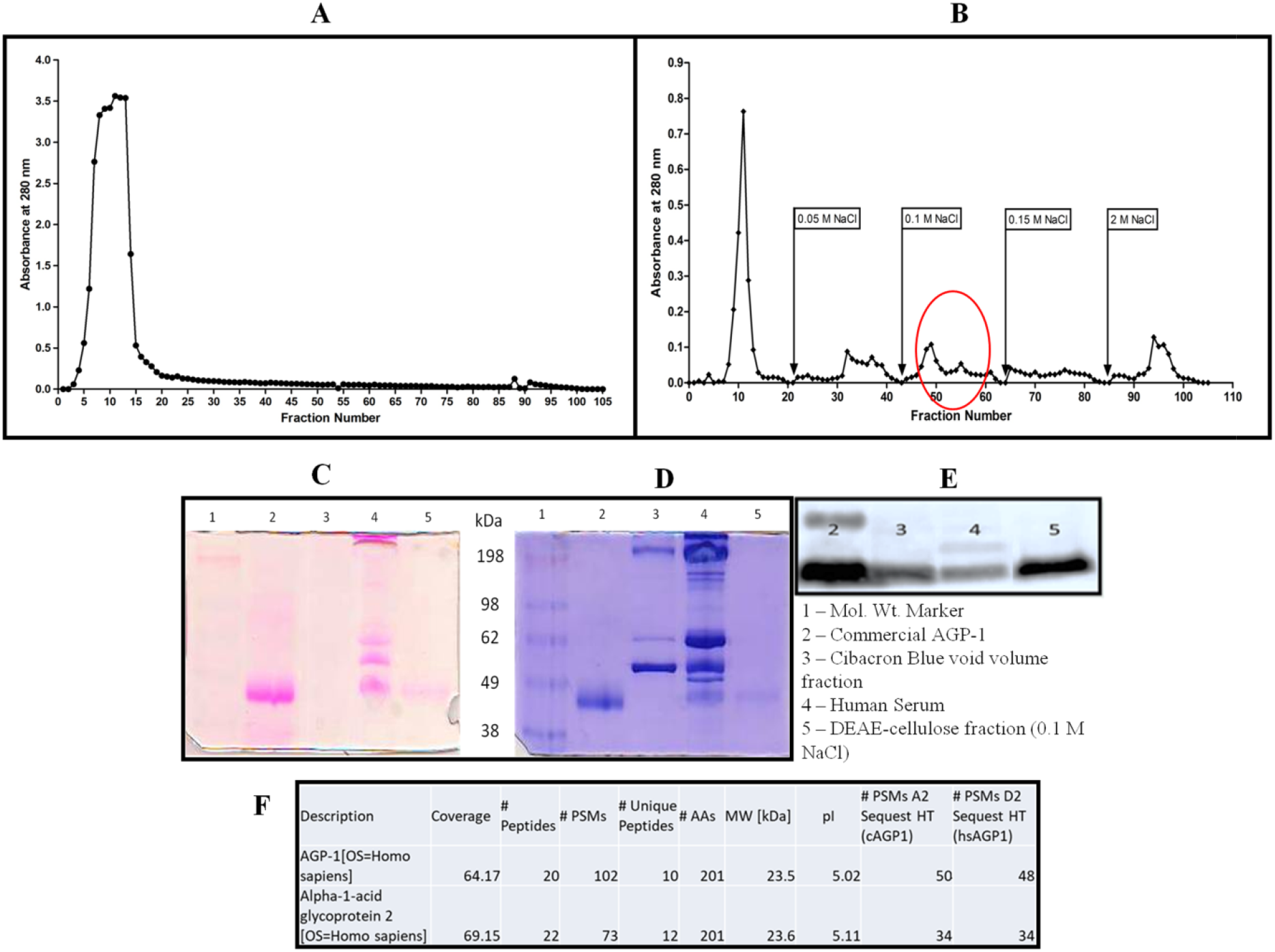
Isolation and characterization of AGP-1 from Human serum. **(A)** Elution profile of human serum proteins on Cibacron blue column. Human serum (2 ml, 80 mg/ml) was loaded on to Cibacron blue pseudo-affinity column and the proteins eluted with 10 mM phosphate buffer pH 7.8 **(B)** Elution profile of concentrated Cibacron blue column void volume fraction from DEAE-cellulose eluted using 30 mM acetate buffer (pH 5.0) containing 0 – 2 M NaCl. The fraction containing AGP-1 is encircled in red. 7.5% SDS-PAGE analysis of AGP-1 after each step of purification was performed and the gel was subjected to both PAS staining **(C)** and Coomassie brilliant blue staining **(D)** and western blot analysis **(E)** using human AGP-1 monoclonal antibody. **(F)** AGP-1 isolated from human serum (hAGP1) and commercially available AGP-1 (cAGP1) was subjected to mass spectral analysis and compared to the AGP-1 protein sequence in the NCBI database. PSMs, AAs, and HT represent peptide spectral matches, amino acids, and hits, respectively.

### AGP-1 inhibits LPS but not BLP – induced MAPK activation

Previous studies have emphasized the MAPK pathway as a critical component of bacterial endotoxin-mediated sepsis (Lu, et al., 2008, Frazier, et al., 2012). We examined the effect of AGP-1 on bacterial endotoxins-mediated MAPK activation in mouse peritoneal macrophages. As expected both LPS and BLP were potent activators of MAPK signaling (Fig 2 B and D). We found AGP-1 alone (25, 50 and 100 μg/ml) activated MAPK signaling and that at the highest concentration that AGP-1 was as effective as LPS in stimulating p38, JNK, and ERK kinase phosphorylation (Fig 2 A and C). Unexpectedly, the combination of AGP-1 and LPS diminished p38 and JNK phosphorylation induced by LPS (100 ng/ml) stimulation in a concentration-dependent manner. Except at the lowest tested concentration (25 μg/ml) where the combination was more effective than LPS alone. AGP-1 also diminished LPS-induced ERK phosphorylation (Fig 2 B and D). In contrast, AGP-1 augmented BLP-mediated p38, JNK, and ERK phosphorylation. Thus, despite the similarity of TLR4 and TLR-2 signaling, AGP-1 elicits opposite outcomes in response to LPS or BLP.

**Figure 2:**
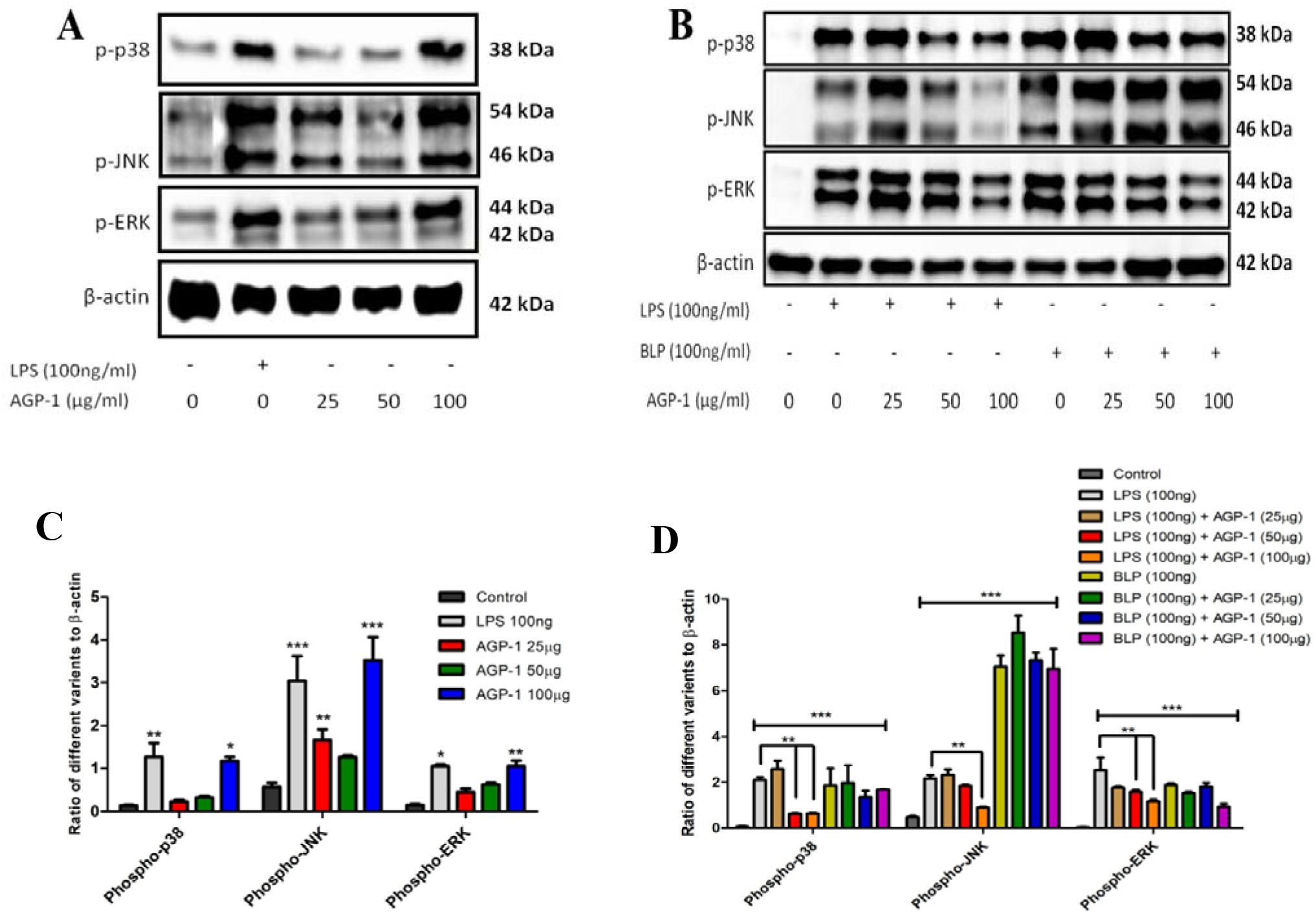
Activation of the MAPK pathway in mouse peritoneal macrophages by AGP-1 and its effect on LPS and BLP induced MAPK pathway. **(A)** AGP-1 affects phospho – p38, JNK and ERK MAP kinases. **(B)** AGP-1 inhibits LPS – induced MAPK pathway activation but not BLP – mediated MAPKs activation. Mouse peritoneal macrophage lysates from the stated amounts of LPS and BLP in the presence and absence of AGP-1 treated cells were prepared using RIPA buffer and immunoblots were developed using specific primary and appropriate secondary antibodies for phospho – forms of MAPKs (p38, JNK and ERK) and β-actin and visualized as described in ‘Methods’. (C) and (D) densitometric analysis of the (A) and (B) blots respectively. Densitometry was done using ImageJ software. The blots are representative of 3 different blots. The data shown are mean ±SEM ***p < 0.001, **p < 0.01 and *p < 0.05 when compared with media, LPS alone and BLP alone.

### AGP-1 selectively augments TLR-2 mediated responses *in vivo*

LPS induced time-dependent mortality in 50% of the injected Swiss albino mice from 16 to 24 hours post-injection (Fig 3A inset). The hepatotoxic D-galactosamine (D-GalN) alone had no effect on the survival of these mice, but greatly enhanced the susceptibility to 15 mg/kg LPS and shortened the time to death, with 100% lethality by 12 hours in response to the combination of D-GalN and LPS (Fig 3 insets). AGP-1 had no effect on mortality by itself but delayed and reduced the lethality of the D-GalN/LPS combination (Fig 3 insets). BLP at 10 or 15 mg/kg did not induce mortality in Swiss albino mice. Addition of D-GalN in combination with the lowest dose of BLP induced mortality over the last hour of the experiment (Fig 3). However, the addition of AGP-1 to the combination of 10 mg/kg BLP and D-GalN advanced the time of lethality to 8 to 9 hours postinjection. AGP-1 itself had no effect on viability. A higher dose of BLP in combination with the hepatotoxin proved to enhance the effect and was more lethal than the lower dose of BLP (Fig 3). Inclusion of AGP-1 enhanced the process where the time to initiation of death was reduced to just 5 hours post-injection (Fig 3). The time to death of multiple animals was not altered by increasing the dose of LPS or addition of AGP-1 (Fig. 3B inset). In contrast, there was a concentration-dependent effect of AGP-1 on the time to BLP/D-GalN induced death, where AGP-1 significantly augmented the time to death at 10 mg/kg BLP but was then unable to modify the rapid death from the highest dose of administered BLP in conjunction with the hepatotoxin. These experiments clearly indicate that AGP-1 selectively augments BLP-mediated, but not LPS-mediated, endotoxemia within a narrow range of insult, highlighting the different response to TLR-4 vs. TLR-2 stimulation.

**Figure 3:**
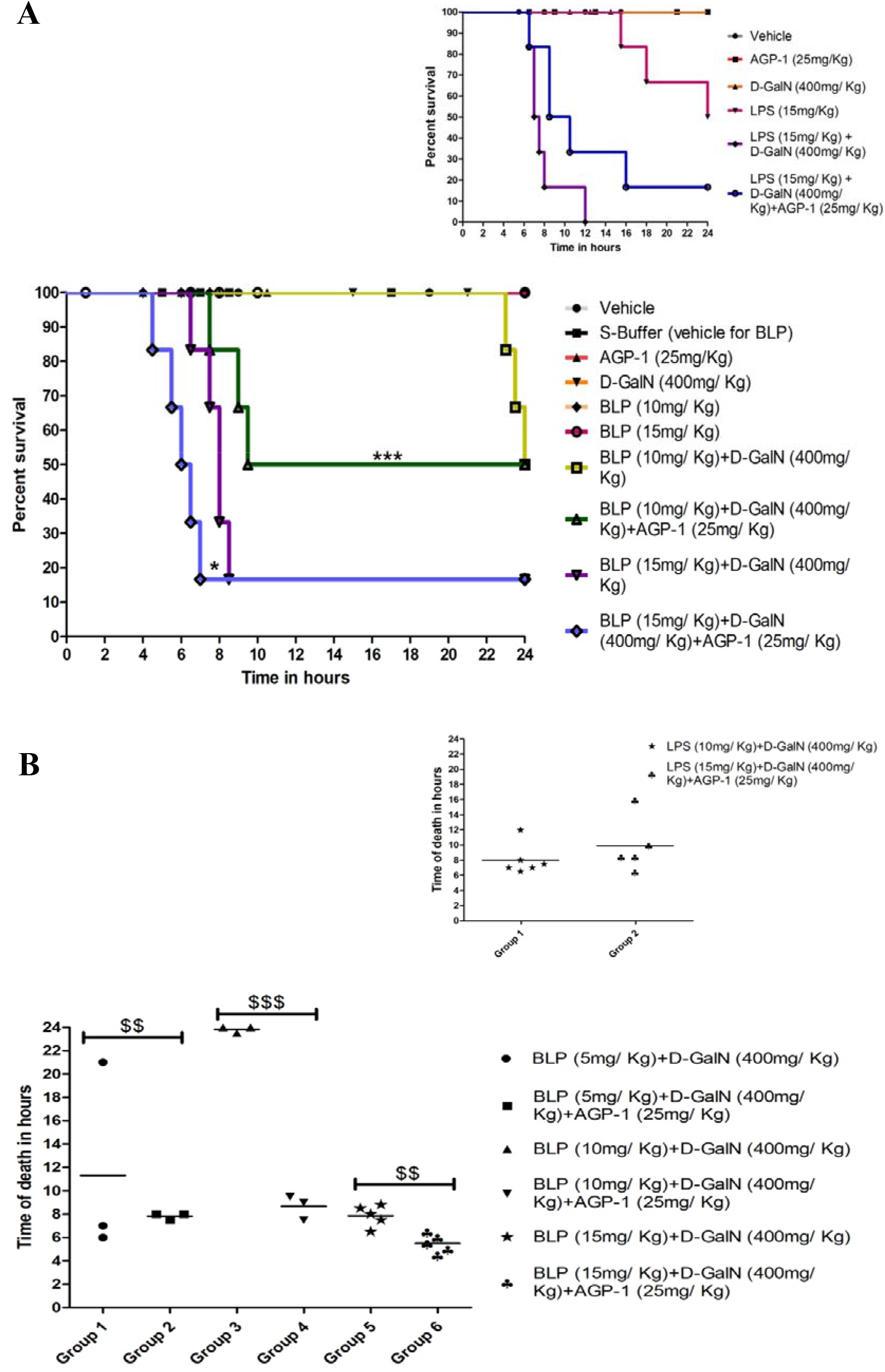
AGP-1 selectively modulates TLR-2 mediated mortality. Swiss albino mice were divided into an indicated number of groups containing six animals each and the mice were injected intraperitoneally with the stated concentrations of BLP and D-GalN in the presence/absence of AGP-1 in a total volume of 500 μl. The survival time was monitored for 24 hours. (A) The graph represents the percentage of survival with respect to time of the groups of mice receiving indicated amounts of BLP and LPS (inset) (15 mg/kg) with or without AGP-1. (B) Time of death of individual animals receiving BLP with or without AGP-1. Inset: LPS (15 mg/kg) was used as a positive control. The result is a representative of three independent experiments. ***p < 0.001 and *p < 0.05 as determined by log-rank test. ^$$$^ p < 0.0001 and ^$$^p < 0.001 as determined by student’s t-test.

### AGP-1 selectively augments TLR-2 mediated neutrophil adhesion *in vitro*

Further, to validated this selective effect of AGP-1 on TLR-2 in humans, we used a primary immune cell, neutrophils as they are the first immune cells to be recruited to the site of inflammation (Kumar and Sharma, 2010). Neutrophils under normal conditions are free-flowing in nature, however, upon inflammation neutrophils adhere to the endothelial layer/extra-cellular matrix via cell adhesion molecules like ß2 integrins, a calcium-dependent pathway (Leick, et al., 2014). This phenomenon was employed to assess the effect of AGP-1 on both TLR-2 and TLR-4 mediated neutrophil adhesion. Labeled neutrophils were stimulated with LPS and BLP with or without AGP-1. AGP-1 alone dose-dependently activated neutrophils. However, neutrophils when incubated with AGP-1 in combination with LPS/ BLP, AGP-1, as seen *in vivo*, preferentially augmented TLR-2 mediated while inhibited the TLR-4 mediated adhesion of neutrophils (Fig 4).

**Figure 4:**
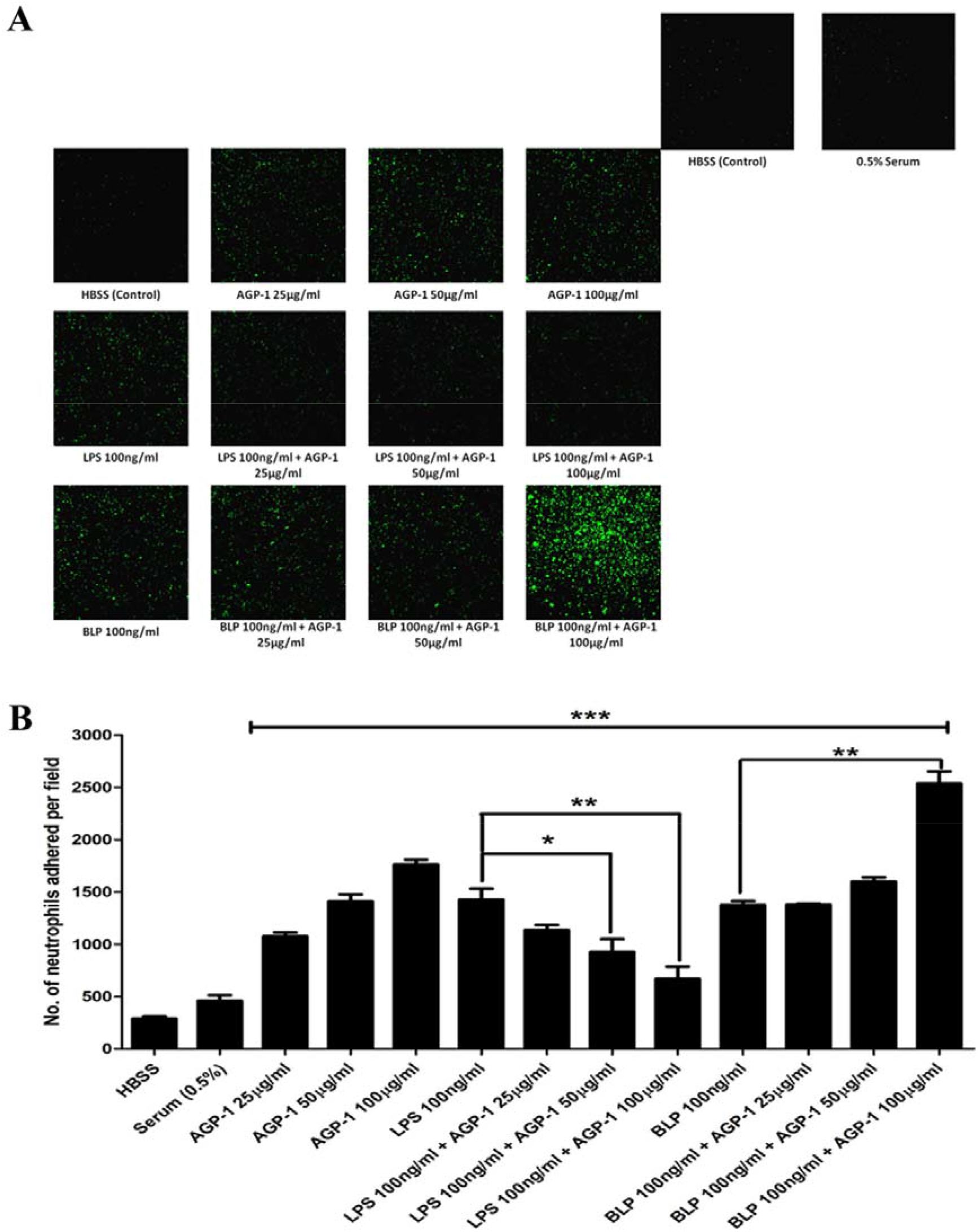
AGP-1 inhibits TLR-4 mediated but potentiates TLR-2 mediated adhesion of neutrophils. **(A)** Neutrophils loaded with calcein-AM in HBSS/A was treated respectively with vehicle, 0.5% serum, LPS/BLP (100□ng/ml) in the presence/absence of AGP-1 (25 – 100 μg/ml). The mixture was incubated for 60□min at 37□°C and the non-adherent PMNs were removed by washing with HBSS. The adherent PMNs were visualized under a fluorescent microscope at a magnification of 40X. Although AGP-1 alone activated neutrophils dose-dependently, it inhibited TLR-4 mediated but potentiated TLR-2 mediated β2-integrin-mediated adhesion of neutrophils to the gelatinous substrate. **(B)** PMN activation was quantified by counting the cells per field using ImageJ software as explained under “Methods”. ***p < 0.0001, **p < 0.001 and *p < 0.01 as determined by ANOVA.

## Discussion

Sepsis is the leading cause of death globally (in ICUs) accounting for 250,000 deaths each year (de Pablo, et al., 2014). Culturable microbes and/ or their products including the membrane components, LPS, and lipoproteins of gram-negative bacteria, mimic the pathobiology of sepsis (Lakshmikanth, et al., 2016, Lakshmikanth, et al., 2016). Although there are several strategies employed to tackle sepsis, more than 100 clinical trials to control sepsis have failed to date (Lakshmikanth, et al., 2016). A new approach is to utilize the endogenous serum components such as the acute phase proteins. However, these proteins exhibit both antiinflammatory (Libert, et al., 1994, Dalli, et al., 2014, Sander, et al., 2010) and pro-inflammatory (Ceciliani, et al., 2002) effects. Dalli et. al. (Dalli, et al., 2014) and Vandevyver et. al. (Vandevyver, et al., 2014) have suggested that the acute phase proteins like alpha-2-macroglobulin (A2MG) can be used as a therapeutic agent against sepsis. With this background in mind, we sought to test the effects of yet another acute phase protein, AGP-1, in a murine model of sepsis, although this protein exhibits both pro- and anti-inflammatory effects (Hochepied, et al., 2003). One of the problems working with AGP-1 is that the presence of multi-molecular forms. AGP-1 is said to have at least 10-12 glyco/ molecular forms in human serum which is mainly dependent on the pathophysiological conditions (Fournier, et al., 2000). De Graaf et. al and Van Dijk et. al showed that there will be an increase in the fucosylation of AGP-1 and the glycoforms expressing the di-antennary glycans during early acute phase response which are dependent on inflammatory cytokines like IL-1, IL-6, and TNF-α (De Graaf, et al., 1993, Van Dijk, et al., 1995). Moreover, there are evidences suggesting that the branching of the glycan part of AGP-1 as well as the degree of fucosylation increases during alcoholic cirrhosis and late-term pregnancy (Biou, et al., 1991, Wieruszeski, et al., 1988). Mackiewicz et. al have suggested that studying AGP-1 glycoforms will help in the diagnosis of cancer (Mackiewicz and Mackiewicz, 1995). We also encountered the problem of the presence of multi-glyco/ molecular forms of AGP-1 in commercial preparations. While studying each of the molecular forms individually would be ideal, isolating them in adequate amounts for experiments would be an insurmountable task. Hence, in this study, we isolated AGP-1 from pooled healthy human serum to apparent homogeneity using fairly simple and effective purification strategies that exhibited uniformity in its molecular form (Fig 1).

We used mouse peritoneal macrophages to test the functionality of this protein in modifying endotoxin-induced MAPK signaling using LPS and BLP to stimulate TLR-4 and TLR-2 respectively since both TLRs are known to activate upstream MAPK pathways to mediate their pro-inflammatory actions (Lu, et al., 2008). Interestingly, AGP-1 alone dose-dependently activated the MAP kinases (p38, JNK, and ERK) (Fig 2 A and C), suggesting that AGP-1 by itself can activate macrophages. Though our results agree with the conclusions of Komori et. al. where AGP-1 induced MAPK signaling in dTHP-1 cells at a much higher concentration of 1 mg/ml accompanied by a retarded activation of MAPKs of 60 – 120 min post-treatment (Komori, et al., 2012), the findings in our study suggest mechanistic underpinnings with purified AGP-1 in primary cells rather than an immortalized cell line. Furthermore, we report for the first time the divergent signaling mechanisms of AGP-1 on the modulation of MAP kinases where the acute-phase protein preferentially inhibited LPS-dependent signaling, but not BLP-induced TLR-2 activation of MAPK (Fig 2 B and D). These results highlight an unexpected diversity of TLR signaling and the complex role(s) of AGP-1 in modifying signaling from PRRs.

Our studies confirm that AGP-1 has a selective effect on murine endotoxemia using models previously established in our laboratory (Lakshmikanth, et al., 2016, Jacob, et al., 2016). We found that AGP-1 at a dose of 25 mg/kg body weight neither potentiated nor protected TLR-4 – mediated mortality (Fig 3). This is in agreement with the previous studies, although surprisingly high doses of both LPS and AGP-1 were used (50 mg/kg and 1 g/kg respectively) (Hochepied, et al., 2003, Hochepied, et al., 2000, Moore, et al., 1997, McCurdy, et al., 2014). In stark contrast, AGP-1 at the same concentration significantly and selectively potentiated TLR-2-mediated mortality (Fig 3), confirming the differential role for AGP-1 as seen in MAPK signaling, where AGP-1 inhibits MAPKs induced by TLR-4, which may, in turn, limit the activation of NF-κB and thereby delays the LPS induced mortality.

In order to check if the murine *in vitro* data can be translated to humans, human neutrophils were stimulated with LPS and BLP with or without AGP-1. Moreover, previous studies have indicated an antiinflammatory role for AGP-1 as it inhibited the migration of neutrophils. Thus, indicating that AGP-1 does not have a role in neutrophil activation (Mestriner, et al., 2007, Matsumoto, et al., 2007, Vasson, et al., 1994). In our study, AGP-1 not only activated neutrophils dose-dependently but was as potent as that of both TLR-2 and TLR-4 agonists (Fig 4). However, neutrophils when stimulated with LPS and/or BLP in combination with AGP-1, it again showed its preferential augmentation of TLR-2 mediated neutrophil adhesion and inhibited TLR-4 mediated effects. Thus validating the results observed *in vivo* murine model of endotoxemia and MAPK signaling in murine macrophages.

In conclusion, we show for the first time that AGP-1 has differential effects on TLR-2 and TLR-4 signaling mechanisms both *in vivo* and *in vitro* that impacts the outcome in terms of survival and lethality in response to specific agonists of these PRRs. The effect of AGP-1 on the TLRs has been summarized in figure 5. Future studies should aim to characterize the physiological functions of different molecular forms of AGP-1 to better understand its role in physiology or pathophysiology.

**Figure 5:**
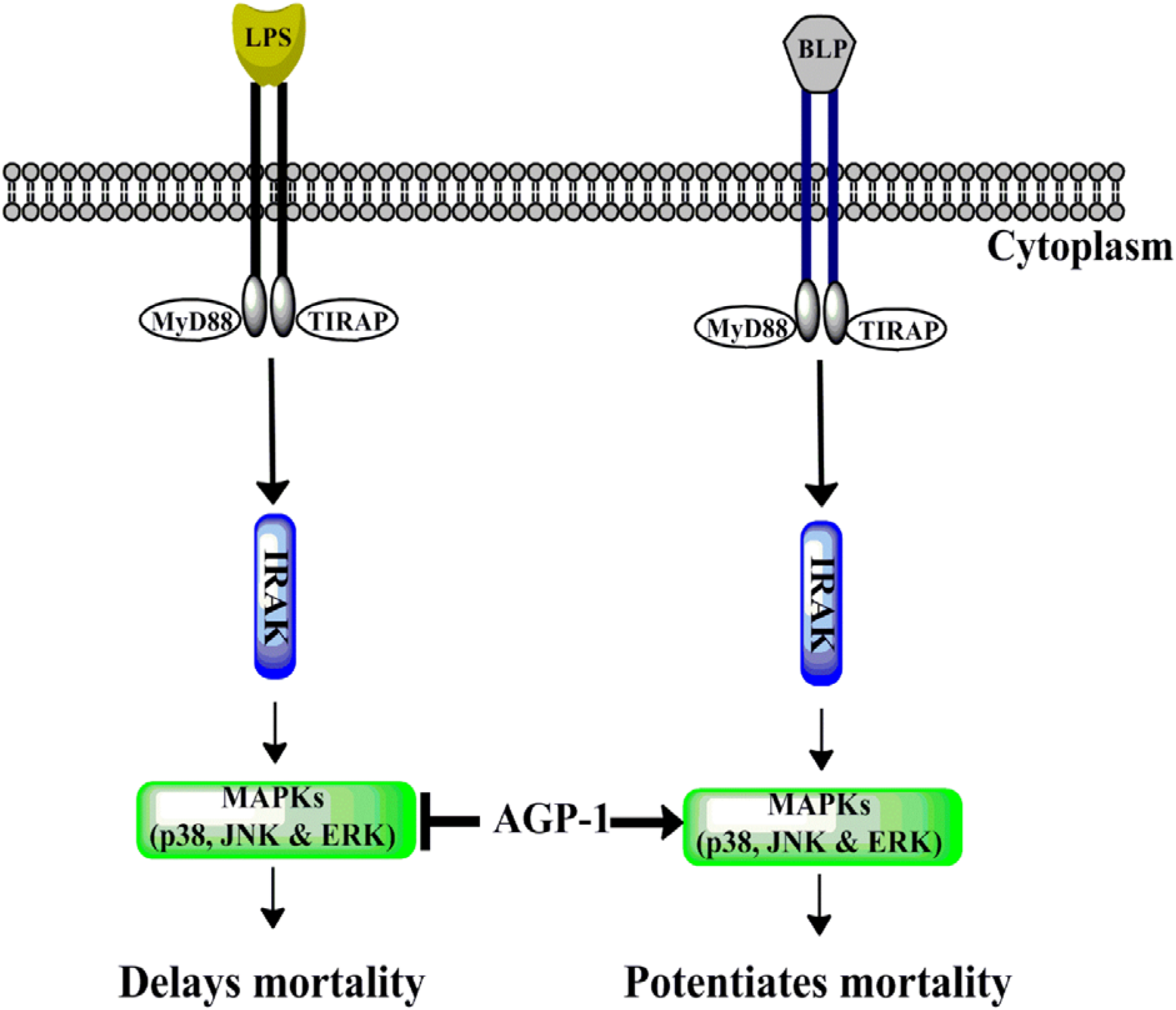
Schematic representation of selective inhibition of MAPKs by AGP-1 induced by LPS but not BLP. AGP-1 selectively inhibits p-38, JNK and ERK MAPKs induced by TLR-4 activation thereby delaying the mortality. However, AGP-1 had no significant effect on the TLR-2 induced MAPK activation but potentiated the BLP mediated mortality.

## Acknowledgments

The authors thank University Grants Commission (UGC), India [for Basic Science Research fellowship – F4-1/2006(BSR)/7-366/2012(BSR) to MSS and F25-1/2014-15(BSR)/7-365/2012(BSR) to VHC, National Fellowship for Higher Education (NFHE-ST) – F1-17/2015-16/NFST-2015-17-ST-KAR-3879/(SA-III/Website) to KVA] and Vision Group of Science & Technology (VGST), Government of Karnataka, India. Authors also thank Dr. Rajesh R, Department of Studies in Molecular biology, University of Mysore, Mysore for generously providing the MAPK antibodies.

## Author Contributions

GKM conceived and designed experiments; MSS, KVA, SPJ, HVC, BW, and VB performed the experiments in the manuscript; GKM, SKP, TMM, BW, and VB analyzed the data; BW and VB contributed reagents/materials/ analysis tools and GKM, SKP, TMM and MSS wrote and edited the manuscript.

